# Quick and efficient approach to develop genomic resources in orphan species: application in *Lavandula angustifolia*

**DOI:** 10.1101/381400

**Authors:** Berline Fopa Fomeju, Dominique Brunel, Aurélie Bérard, Jean-Baptiste Rivoal, Philippe Gallois, Marie-Christine Le Paslier, Jean-Pierre Bouverat-Bernier

**Affiliations:** iteipmai, Melay BP 80 009, 49120 Chemillé-en-Anjou, France; INRA Institut National de la Recherche Agronomique, US1279 Etude du Polymorphisme des génomes Végétaux, CEA-IG/CNG Centre National de Génotypage, 2 rue Gaston Crémieux, 91057 Evry, France; CRIEPPAM, Les Quintrands, Route de Volx, 04100 Manosque, France

## Abstract

Next-Generation Sequencing (NGS) technologies, by reducing the cost and increasing the throughput of sequencing, have opened doors of research efforts to generate genomic data to a range of previously poorly studied species. In this study, we proposed a method for the rapid development of a large scale molecular resources for orphan species. We studied as an example *Lavandula angustifolia*, a perennial sub-shrub plant native from the Mediterranean region and whose essential oil have numerous applications in cosmetics, pharmaceuticals, and alternative medicines.

We first built a ‘Maillette’ reference Unigene, compound of coding sequences, thanks to *de novo* RNA-seq assembly. Then, we reconstructed the complete genes sequences (with exons and introns) using a transcriptome-guided DNA-seq assembly approach in order to maximize the possibilities of finding polymorphism between genetically close individuals. Finally, we used these resources for SNP mining within a collection of 16 lavender clones and tested the SNP within the scope of a phylogeny analysis. We obtained a cleaned reference of 8, 030 functionally annotated ‘genes’ (*in silico* annotation). We found up to 400K polymorphic sites, depending on the genotype analyzed, and observed a high SNP frequency (mean of 1 SNP per 90 bp) and a high level of heterozygosity (more than 60% of heterozygous SNP per genotype). We found similar genetic distances between pairs of clones, related to the out-crossing nature of the species, the restricted area of cultivation and the clonal propagation of the varieties.

The method propose is transferable to other orphan species, requires little bioinformatics resources and can be realized within a year. This is the first reported large-scale SNP development on Lavandula angustifolia. All this data provides a rich pool of molecular resource to explore and exploit biodiversity in breeding programs.

## Introduction

Next-Generation Sequencing (NGS) technologies, by reducing the cost and increasing the throughput of genotyping and sequencing, have opened doors of research efforts to generate genomic data to a wide range of species that had not yet benefited. In particular, reduced-representation sequencing methods made it possible to develop markers and genomic sequences without any prior genomic information as a reference genome, and thus are particularly suitable to these so-called ‘orphan’ (or ‘minor’) species [1,2]. These methods are either based on the sequencing of DNA fragments after use of restriction enzymes (Restriction-site Associated DNA sequencing (RAD-seq), Genotyping-By-Sequencing (GBS), to cite few), or on the sequencing of the expressed fraction of the genome (exome sequencing, RNA sequencing (RNA-seq)) [3,4].

It is relevant to take advantage of this technological leap to work on these species ‘neglected’ by the global academic research, but with strong local economic importance [5]. In addition, it is increasingly recognized that a better exploitation of biodiversity in agriculture would help to face the new issues (diseases, droughts) related to climate change [6–8].

In this study, we proposed a methodology for the development, within a year, of a large scale molecular resources for an orphan species. We studied as an example *Lavandula angustifolia*. Lavender is a perennial sub-shrub plant native from the Mediterranean region and is best known for its essential oils that have numerous applications in perfume, cosmetics, pharmaceuticals, and alternative medicines [9–11]. Lavender is one major species in the Medicinal and Aromatic Plants (MAP) field: the lavender culture (which includes true lavender, *L.angustifolia*, and a related species, the lavandin (*Lavandula x intermedia*)) represents 43% of the surface area of MAPs in France [12]. Another important specificity of lavender is its ability to produce yields under water-limited environments and on stony mountain soils where no other successful farming is possible [10]. Moreover, the cultivation of a limited number of varieties has drastically narrowed the genetic diversity used for breeding programs: three varieties represent 50% of lavender cultivated area [13] and the clone “Grosso” represents 80% of lavandin cultivated area. Developing genomic tools for the lavender will allow to explore a broad range of genetic diversity available in the species and its relatives and to provide new varieties improved for traits of agronomic interest.

To date, few molecular resources have been developed on *Lavandula angustifolia* and no SNP markers have been reported. As of June 2017, 204 nucleotide sequences had been deposited in the NCBI’s GenBank database for *Lavandula* species, with more than 50% being sequences of *Lavandula angustifolia*. Most of them (~25%) are related to synthesis or regulation of essential oils [14–21]; or are isolated from the chloroplast genome (~30%). Moreover, EST-derived SSR markers have been recently developed and successfully tested for transferability between *Lavandula* species [22]. In this study, we firstly developed genomic reference sequences thanks to *de novo* RNA-seq assembly and transcriptome guided DNA-seq assembly. Secondly, we used these resources for SNP mining and finally tested the SNP within the scope of a phylogeny analysis of lavender clones.

We selected SNP markers among other molecular markers because they are stable, simple to detect and easily amenable to a high-throughput automation. In addition to their ‘technical’ advantages, SNPs, due to their abundance at genome-wide level, are the most desirable, precise and efficient markers for developing high-density genome scans[23–26].

To develop genomic sequences, we chose the *de novo* RNA-seq assembly among various available methods because it allows (i) to develop a reservoir of reference DNA sequences that can be used for subsequent analyzes and (ii) to functionally annotate the *de novo* assembled sequences thanks to comparative genomics and thus to have access to candidate genes for breeding purpose. We then included introns in our reference sequences through a pipeline developed by Aluome *et al*. [27], increasing the opportunity of SNP detection in case of low genetic distance between individuals.

The results presented herein offer a solid base for the initiation of population genetics studies, DNA fingerprinting and the development of efficient genomic-assisted selection strategies for *Lavandula angustifolia* and the related species of the *Lamiaceae* family.

## Material and methods

### Plant material

The plant material was selected to be representative of the phenotypic variation (morphological, essential oils content) observed in the lavender clones cultivated nowadays (S1 Table). The lavender selection includes two putative geographical origins: 2 clones named B6 and B7 known for their Bulgarian origin; and 13 clones from the south-east of France, mainly from Albion plateau. Almost all the clones were collected from fields of open-pollinated varieties and it was difficult to trace the exact origin and pedigree of each clone. However, some information was provided by the CRIEPPAM (Regional Interprofessional Center for Experimentation in Aromatic and Medicinal Fragrant Plants, Manosque, France) and is indicated in S1 Table. The clone‘Maillette’ was chosen to construct the reference sequence of lavender because of its large use in the lavender production area. The lavandin ‘Grosso’, a natural sterile interspecific hybrid between *Lavandula angustifolia* and *Lavandula latifolia*, was also included because of it high economic importance in the MAP sector.

All the clones were provided by the Technical and Interprofessional Institute of Perfume and Medicinal Plants (ITEIPMAI - Chemillé, France) and were maintained at the ARDEMA (arid mountain research and development association) experimental farm (Mévouillon, France).

## RNA and DNA extraction, Library construction and Sequencing

### RNA and DNA extraction

To isolate RNA, samples were collected at the end of May. Due to the early date of collection leaves and root samples were collected from an adult plant of the clone ‘Maillette’ maintained in the field and flower buds were collected from an adult plant maintained in a green house. Collected samples were immediately frozen and conserved in liquid nitrogen until use. The samples were grounded in liquid nitrogen. Total RNA was isolated for each tissue sample using Plant RNA Isolation Mini Kit (Agilent, Santa Clara, CA, USA), according to manufacturer’s instructions.

To isolate DNA, leaves samples were collected from 15 lavender clones (including Maillette) and the lavandin ‘Grosso’ at an adult stage, in the field, and immediately frozen and stored in dry ice until use. Total DNA was isolated independently from leaves of the 16 clones according to manufacturer’s recommendations using the NucleoSpin Plant II kit (Macherey-Nagel, Düren, Germany).

After isolation, the yield, purity and integrity of RNA and DNA samples were analyzed using a Molecular Devices Multimode Plate Reader SpectraMax M3 (Molecular Device) and a TapeStation (ADN) or a BioAnalyzer (ARN) (Agilent) device.

### Library Preparation and Sequencing

Libraries were prepared independently for each tissue sampled.

After the total RNA was extracted, the cDNA stranded libraries were prepared using the TruSeq stranded mRNA Sample Preparation Kit (Illumina Inc., San Diego, CA, USA) according to manufacturer’s recommendations. The DNA libraries were prepared with the TruSeq DNA PCR-Free Sample Preparation kit (Illumina Inc., San Diego, CA, USA) according manufacturer’s recommendations.

The RNA and DNA libraries were paired-end sequenced on an Illumina HiSeq 2500 (2*150pb) at the French National Center for Genotyping (Evry, France).

All sequencing reads were deposited into the Short Read Archive (SRA) of the National Centre for Biotechnology Information (NCBI) and can be accessed under the Bioproject number PRJNA391145.

## Data Analysis

### DNA-seq and RNA-seq trimming

The raw paired-end reads produced following RNA and DNA sequencing were filtered with CLC Genomics Workbench 8.5 (https://www.qiagenbioinformatics.com/), hereafter named CLC, to obtain high-quality cleaned reads. The Illumina adapter sequences, low quality sequences (limit = 0.001), ambiguous nucleotides (no “N” allowed), and short sequences (minimum length = 70 nucleotides) were removed during the trimming process.

### *De novo* assembly of leaf, flower bud and root transcriptome of Maillette

The pipeline used to build our lavender reference Unigene from RNA-seq data is presented on Fig 1. The currently popular tools used for the *de novo* RNA-seq assembly are all based on the construction of a *De Bruijn* graph. However, each tool has its advantages and its limits and produce different types of bioinformatically derived artefacts. In a recent study, Cerveau and Jackson [28] have shown the interest of surveying the outputs of different assembly tools to generate a high-quality transcriptome. In the present study, we chose to perform de novo assemblies with CLC v 8.5 and TRINITY v 2.1.1[29,30]. The cleaned paired-end reads from RNA sequencing of leaf, flower bud and root of the lavender clone ‘Maillette’ were pooled and used for the *de novo* transcriptome assemblies. Since reads 1 were of better quality than read 2, they were used to build the primary *De Bruijn* graph and reads 2 were used to resolve bubbles in the graph. The k-mer size values used were 64 bases for CLC and 25 bases for TRINITY (fixed default value).

**Fig. 1:**
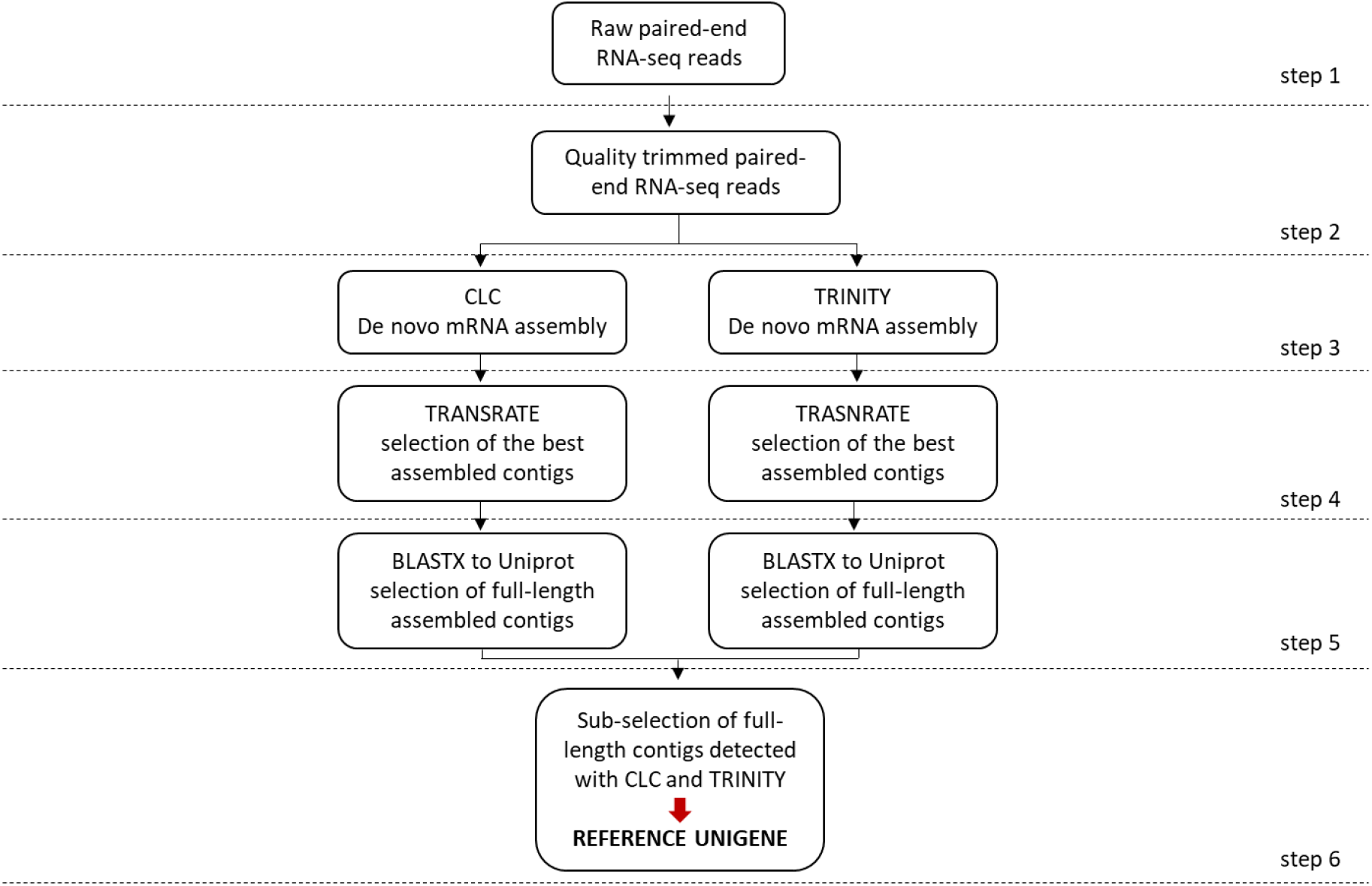
RNA-seq assembly for the construction of the reference Unigene pipeline.

The quality of the raw assemblies was assessed with TRANSRATE [31]. TRANSRATE maps paired-end reads back to the raw contigs and calculate metrics (percentage of reads mapping back to contigs in proper pairs, number of contigs with a predicted open reading frame, estimation of fragmented transcripts…), in order to assess how well the assembled contigs are supported by sequencing data. On this basis, a score is determined for each contig and the set of contigs with best scores are gathered into an “optimized assembly”. TRANSRATE also computes a global score that allows comparing two or more assemblies performed with the same initial dataset but with different tools or settings.

To evaluate the completeness of the assemblies, we performed a BLASTx+ (e-value 1e-20) alignment of the optimized assemblies against SwissProt database (http://www.uniprot.org/) using a script developed by the TRINITY developer’s team (https://github.com/trinityrnaseq/trinityrnaseq/wiki/Counting-Full-Length-Trinity-Transcripts). This analysis allowed to determine (i) the number of unique best blast hit (BBH) found with contigs from each optimized assembly and (ii) the percentage of coverage of the hit sequence by the contig.

We finally took advantage of both assemblies to build the reference Unigene. This Unigene is compound of the sequences assembled with TRINITY or CLC that covered more than 70% of their BBH in SwissProt database with more than 30% identity. In case of redundancy within an assembly (*i.e*, several contigs matched the same hit in the database), we only kept the longest contig in the Unigene. Likewise, when two sequences built by CLC and TRINITY had the same BBH, we only kept the longest form in the Unigene or, in cases of equality, the one with the highest alignment similarity percentage.

### *In silico* functional annotation of the reference Unigene

To further describe biological functions related to the Unigene sequences, we ran the Trinotate pipeline (https://trinotate.github.io/) which is a comprehensive annotation suite adapted to *in silico* annotation of *de novo* assembled transcriptome. The analysis includes homology searches (evalue 1e-5) to SwissProt database, protein domain identification using HMMER and Pfam databases, protein signal peptide and transmembrane domain prediction with Signalp and tmHMM servers, and search for Gene Ontology (GO) terms and KEGG pathways [32,33]. Insofar as the ability to annotate sequences based on similarity search against databases depends on the completeness of these databases, we completed the annotations using supplementary databases and tools. The online tools KAAS (http://www.genome.jp/kegg/kaas/) [53] was used with default parameters to decipher associated KEGG pathways by aligning Unigene sequences against the KEGG GENES database. The online tool TRAPID (http://bioinformatics.psb.ugent.be/webtools/trapid/) [54] was also used with default parameters to discover associated GO terms. TRAPID performs a similarity search against implemented databases (OrthoMCL-DB version 5 and PLAZA 2.5), and an open-reading frame detection. Results are then combined to identify coding sequences, assign transcripts to gene families and generate GO annotation.

### Iterative targeted assembly of lavender genes

The Unigene sequences and the reads from the DNA sequencing of the clone ‘Maillette’ were used to performed iterative mapping /assembly steps to build a set of reference gene sequences (with exons and introns) as described in Aluome *et al* [27] (Fig. 2, S9 File), hereafter called Genespace.

**Fig. 2:**
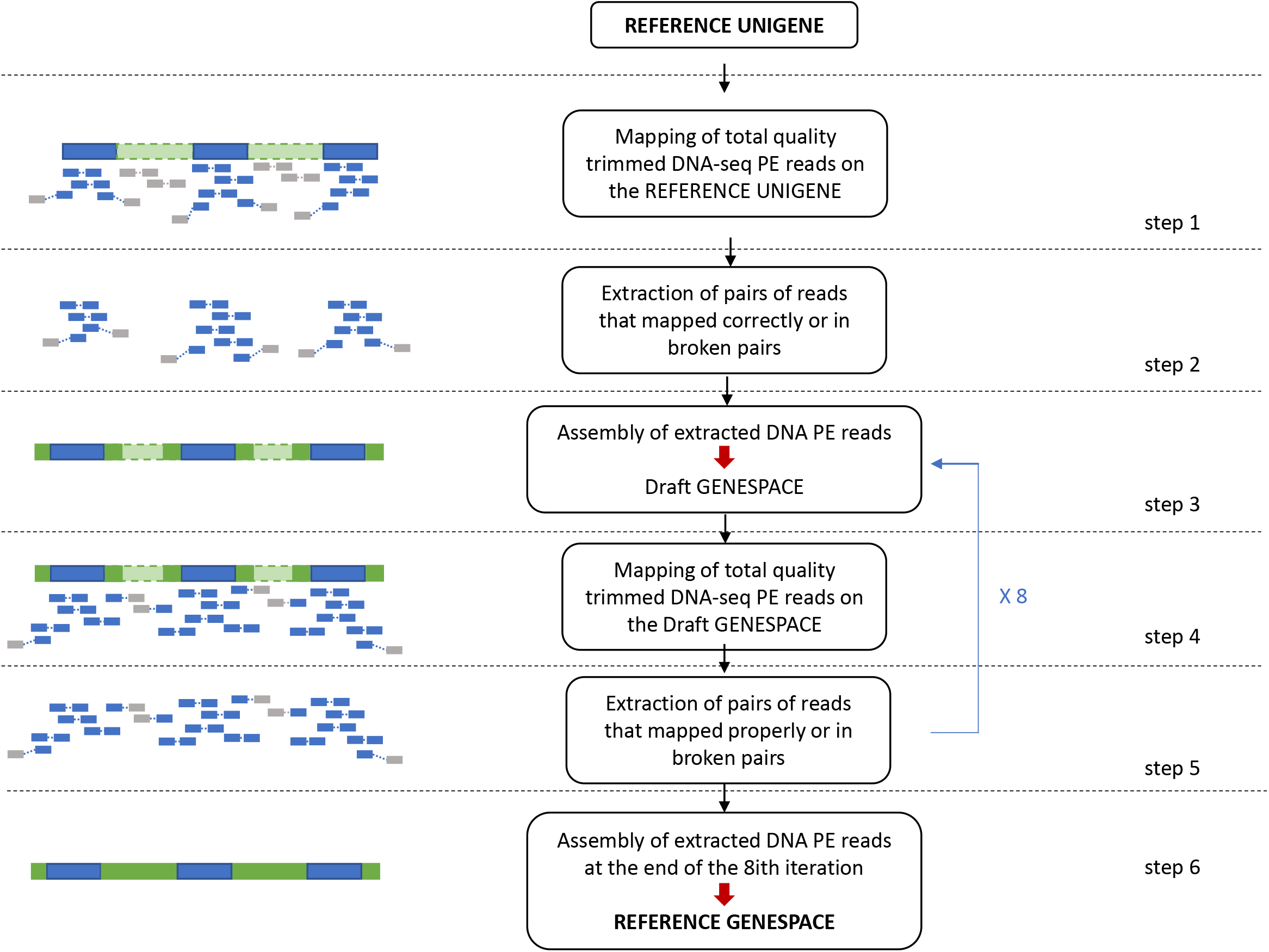
Transcriptome-guided DNA assembly for the construction of the reference Genespace pipeline. Left diagram: blue boxes are gene exons and green boxes are gene introns. Dashed lines indicate missing sequences. Paired-end DNA-seq reads are blue when both member of the pair map on the reference sequence or blue and grey when they map in broken pair.

Briefly, the first iteration consists in mapping the whole-genome DNAseq reads on the reference sequences of the Unigene (step 1, Fig. 2). Then, the reads that mapped in pairs and broken pairs (*i.e*, only one member of the pair mapped) are extracted (step 2, Fig. 2) and assembled into *de novo* contigs (step 3, Fig. 2). For further iterations, the total reads from DNA sequencing are mapped on the new sequences assembled at the end of the (*i-1*)^th^ iteration (repeat of steps 3 to 5, Fig. 2). We stopped the iterations when the maximum number of sequences from the reference Unigene were re-built in the draft Genespace (in our case, at the 8i*th* iteration) thus getting our reference Genespace (step 6, Fig. 2).

CLC Assembly Cell (https://www.qiagenbioinformatics.com/) v 5.0 was used for reads mapping (CLC_ref_assemble_long tool) and read extraction (sub_assembly tool) steps. Mapping was less stringent at the first iteration (step 1, Fig. 2: 100% of the read length with a similarity percent of 95%) than for other iterations (step 4, Fig. 2: 100% of the read length with a similarity percent of 98 %) in order to take into account exons/introns junctions at the first step of mapping. For assembly steps, we used idba_ud v. 1.0.9 software [34]. This tool offers the advantage to use the new sequences assembled at the iteration *i-1* to ‘guide’ the assembly at the iteration *i* (--input-long-read option) and thus allowing to keep the information build at the iteration *i-1* while assembling new sequences at iteration *i*. Parameters used for idba_ud for all iterations were a minimum kmer size of 25 nucleotides and a maximum kmer size of 100 nt (--mink 25 --maxk 100), with an increment of 5 nucleotides (--step 5) and a similarity value of 1 nucleotides (--similar 1).

To assess the reconstruction of the Unigene sequences with a successful intron introduction, we performed a BLASTN alignment (e-value 1e-6) of Unigene sequences against the sequences built at the end of each iteration of DNA assembly.

### SNP discovery

Filtered DNA-seq paired-end reads of the 15 lavender clones and of the lavandin ‘Grosso’ were mapped against the Genespace sequences with CLC software. To be included in the mapping, at least 95 % (90% for ‘Grosso’) of the read (length fraction = 0.95 or 0.90) must be aligned to the reference sequence with at least 90% identity (80% for ‘Grosso’) (similarity fraction =0.90 ou 0.80). Moreover, reads with non-specific matches (*i.e*. reads mapping equally well at several alignment positions) were excluded from the mapping.

SNP detection was performed with CLC and GATK v 3.6 [35–37] tools. It has been demonstrated that a significant improvement of SNP calling –in terms of further successful genotyping of the SNP-can be obtained by focusing on SNP discovered by more than one method [38]. According to the initial sequencing depth of the genotypes studied; we applied a maximal depth cutoff of 50X to consider a site for SNP discovery. This maximum cutoff was applied to prevent SNP detection in regions with very high depth of coverage that could correspond to repeated DNA regions.

For variant discovery with CLC we used the *Fixed Ploidy Level* tool. A diploid model for SNP calling (ploidy level = 2) was selected with a minimum read coverage of 5, a maximum read coverage of 50, a minimum non-reference allele count of 2 with a minimum frequency of 40% (calculated as “number of non-reference alleles at the site”/ “total site coverage”). Broken pairs and non-specific reads were not used for SNP detection. The minimum base quality required for the putative SNP and the 5 bases on both sides of the SNP was of 25. For variant detection with GATK, mapping files generated with CLC were exported in *bam* format files to be used in GATK. We used HaplotypeCaller, SelectVariants and VariantFiltration tools included in GATK to perform the analyses. A maximum read coverage of 50 and a minimum per base quality of 30 were required to consider a polymorphic site as a putative variant (since, with GATK, we could not provide a quality parameter for the bases in the vicinity of the called SNP, we defined a quality threshold for the called SNP higher in GATK than in CLC). We then applied SelectVariants and VariantFiltration programs to only select SNP in each vcf files and to apply filters to SNP calling similar to those applied with CLC. Finally, in order to prevent from spurious SNP calling, we applied SelectVariant tool to extract the SNP that were concordant (position and genotype information) between CLC and GATK vcf output files to get a cleaned set of putative SNPs.

### Assessment of the lavenders genetic relationship

From the filtered putative SNPs set, we selected SNPs that generated no missing genotype data. The analyses were based on the ‘Population Structure’ workflow from the Grundwald laboratory (https://grunwaldlab.github.io/Population_Genetics_in_R/Pop_Structure.html). The R packages ‘poppr’ version 2.7.1[39,40], ‘adegenet’ [41,42] and ‘ade4’ [43] were used to calculate absolute genetic distances with ‘Prevosti’s method’ [44] (a method suited to SNP data), to perform a principal component analysis (PCA) and built an Neighbor joining phylogenetic tree with 1000 bootstrap.

## Results

### RNA-seq and DNA-seq trimming

A total of 169,082,890 (84,541,445 pairs) of RNA paired-end (PE) reads were sequenced reaching 25.5 Gigabases (Gb) (S2 Table) from leaves, roots and flower buds of ‘Maillette’. After trimming on quality, length, ambiguous nucleotide and adapters, a total of 144,322,837 reads totaling 19.7 Gb (77% of initial dataset) remained for *de novo* transcriptome assembly with a mean PHRED score per read around 40. Depending on the genotype, the number of paired-end DNA reads sequenced ranged from 29,985,684 to 145,227,504(S2 Table). Given that the genome size of *Lavandula angustifolia* is approximately 1Gb (Iteipmai CASDAR 2011, [45]), this corresponds to a theoretical mean depth of coverage ranging from 5X to 22X. The clone Maillette reaches the highest sequencing depth because it was used for the transcriptome-guided DNA assembly described below.

### De novo transcriptome assembly

Results of *de novo* assemblies are presented on Table 1. On overall, TRINITY generated more contigs of a longer size than CLC. A total of 187,584 sequences (109.6 Mb) were assembled with CLC and 280,062 (220.3 Mb) with TRINITY The size of the contigs ranged from 200 bp to approximately 15 kb with both methods and with mean contig sizes of 584.68 bp (CLC assembly) and 786.78 bp (TRINITY assembly). TRANSRATE uses reads data in pairs to assess the quality of assemblies (Table 1). A total of 66,107,704 paired-end reads (92% of cleaned reads) remained in pairs after independent trimming of the reads 1 and the reads 2. More than 62 million of pairs mapped back to the contigs assembled with TRINITY and CLC. Of these mapped reads, almost 60 % were assessed to be mapped in “good” pairs according to the TRANSRATE standards, meaning for which both members of a pair were aligned on the same contig, in the correct orientation and without overlapping either end of a contig. If the TRANSRATE score obtained for CLC raw assembly (0.154) was better than for TRINITY raw assembly (0.108), the scores were similar for optimized assemblies with a score of 0.184 and 0.198 for CLC and TRINITY assemblies, respectively.

**Table 1:** Assembly metrics for Maillette *de novo* transcriptome assembly with CLC and Trinity tools.

| Description | Maillette TRINITY | Maillette CLC |
| --- | --- | --- |
|  | Assembly metrics |  |
| Number of assembled contigs | 280,062 | 187,584 |
| Minimum contig length (bp) | 201 | 200 |
| Maximum contig length (bp) | 15,6 | 14,6 |
| Assembly size (bp) | 22,0348,569 | 109,677,226 |
| Mean contig length (bp) | 786.78 | 584.68 |
| Number of contigs > 1 kb | 70,988 | 25,762 |
| Number of contigs > 10 kb | 20 | 1 |
| Number of contig with predicted ORF (percent) | 96,630 (66.20%) | 50,399 (72.47%) |
| N50 | 1250 | 739 |
| GC percent | 42.84 | 42.97 |
| Linguistic complexity | 0.14942 | 0.11786 |
|  | TRANSRATE quality assessment |  |
| Number of available PE read | 66,107,230 | 66,107,230 |
| Number of mapped PE reads (percent) | 62,126,811 (94 %) | 62,068,704 (94 %) |
| good_mappings (percent of mapped reads) | 37,495,608 (60 %) | 35,932,087 (58 %) |
| potential_bridges | 91,104 | 57,505 |
| Number of base uncovered (percent) | 15,251,707 (7 %) | 507410 (0.5 %) |
| Number of contigs with at least 1 uncovered base (percent) | 152,145 (54 %) | 58,033 (31 %) |
| Contings with a mean per-base coverage <1 (percent) | 11,086 (3.95 %) | 212 (0.11 %) |
| Contings with a mean per-base coverage < 10 (percent) | 168,024 (59.99%) | 73,318 (39.08 %) |
| Number of contigs putatively segmented (percent) | 30,662 (10.95 %) | 23,851 (12.72 %) |
| Raw assembly score | 0.10897 | 0.15453 |
| Optimal assembly score | 0.19831 | 0.18401 |
|  | Completeness assessment with BLASTX alignment |  |
| Total number of optimised contigs | 194,706 | 147,672 |
| Number of contigs with BBH (percent) | 78,929 (40.53%) | 51,821 (35.09%) |
| Number of unique BBH | 18,024 | 16,270 |
| Number of hit with >= 70% coverage | 10,508 | 8,286 |
| Mean contigs (or HSP)/ BBH | 4.37 | 3.18 |

### Finally, a pool of 147,672 contigs from CLC raw assembly and 194,706 contigs from TRINITY raw assembly were selected as two optimized assemblies for downstream analyses

To assess the completeness of the optimized contigs sets, we used a TRINITY utility (see Methods), to perform a BLASTX alignment against SwissProt database (e-value 1e-20) (Table 1). From the optimized contigs sets, a total of 51,821 (35%) contigs built with CLC and 78,929 (40.5%) contigs built with TRINITY had a hit in the database. There was redundancy in both optimized assemblies since the 51,821 sequences from CLC optimized assembly matched 16,270 unique best blast hit (BBH) and the 78,929 sequences from TRINITY optimized assembly matched 18,024 unique BBH. This result suggested the presence of homologous sequences such as gene families in the assemblies. **Thus, we took advantage of both assemblies** (see Methods) **and used the BLASTX results to construct a non-redundant reference Unigene of 10,060 contigs**. From those, 8,951 (89%) contigs that were deciphered with CLC and TRINITY, 739 contigs (7%) were specifically assembled with TRINITY and 370 contigs (4%) were built only with CLC. Selected sequences have length ranging from 202 bp to 15,595 bp and a median value of 1,480.5 bp (Fig 3B).

### *In silico* annotation of the reference Unigene

The reference Unigene was annotated with various pipeline and software to retrieve gene ontology (Trinotate, TRAPID), KEGG pathways (Trinotate, KAAS), gene families (TRAPID) and additional functional annotation (Trinotate pipeline). Detailed results from these softwares are summarized in S3 Table. Out of the 10,060 sequence of the reference Unigene, 9,816 (97.6%) had a BLASTP hit against swissprot database, 8,928 (88.7%) had at least one GO annotation, 4,931(49%) had at least one KEGG pathway annotation and 4,647 (46%) had a KEGG and a GO annotation. Search of conserved domain against Pfam database allowed to increase the rate of sequence annotated with a GO term or a KEGG ontology (S3 Table). The identification of various GO terms and KEGG pathways indicated that we assembled sequences related to a relatively diversified panel of protein functions.

### Introns insertions in the reference Unigene sequences

According to Aluome *et al*.[27], an iterative transcriptome-guided DNA assembly was performed to insert introns into Unigene sequences (Fig. 2).

Out of the 10,060 sequences of the reference Unigene, 8,147 (81%) sequences were found in the DNA sequences assembled at the end of the 8th iteration (Fig 3A). From those 8,147 sequences, 3,556 (43.64%) matched longer sequences in the DNA assembly than in the Unigene; 1,164 (14.28%) were concatenated in 550 sequences with the DNA assembly; and 3,427 (42.06%) sequences matched shorter sequences in the DNA assembly. A total of 1,913 sequences of the Unigene (19%) did not have a hit in the assembled DNA-seq and were later identified as being contaminant sequences (see next section). The alignment (BLASTN) of the Unigene contigs against their corresponding “gene sequence” from the DNA-seq assembly allowed to estimate the number and the length of introns and exons in the “gene sequences” (Fig. 4). Thus, the number of exon per sequence ranged from 1 to 24 with a mean value of 3 (median =2); and the number of introns per sequence ranged from 0 to 23 with a mean value of 2 (median =1). In addition, we noticed that the size of the exons and introns was roughly equivalent. Exons size ranged from 54 bp to 4,000 bp (mean=288 bp, median=189 bp), and introns size ranged from 0 to 3,800 bp (mean=235 bp; median=114 bp). The mean percent of identity for an HSP alignment was of 98 %, indicating that, when existing, sequences reconstructed with the DNA-seq assembly were highly similar to the ones built with de novo RNA-seq assemblies.

**Fig. 3:**
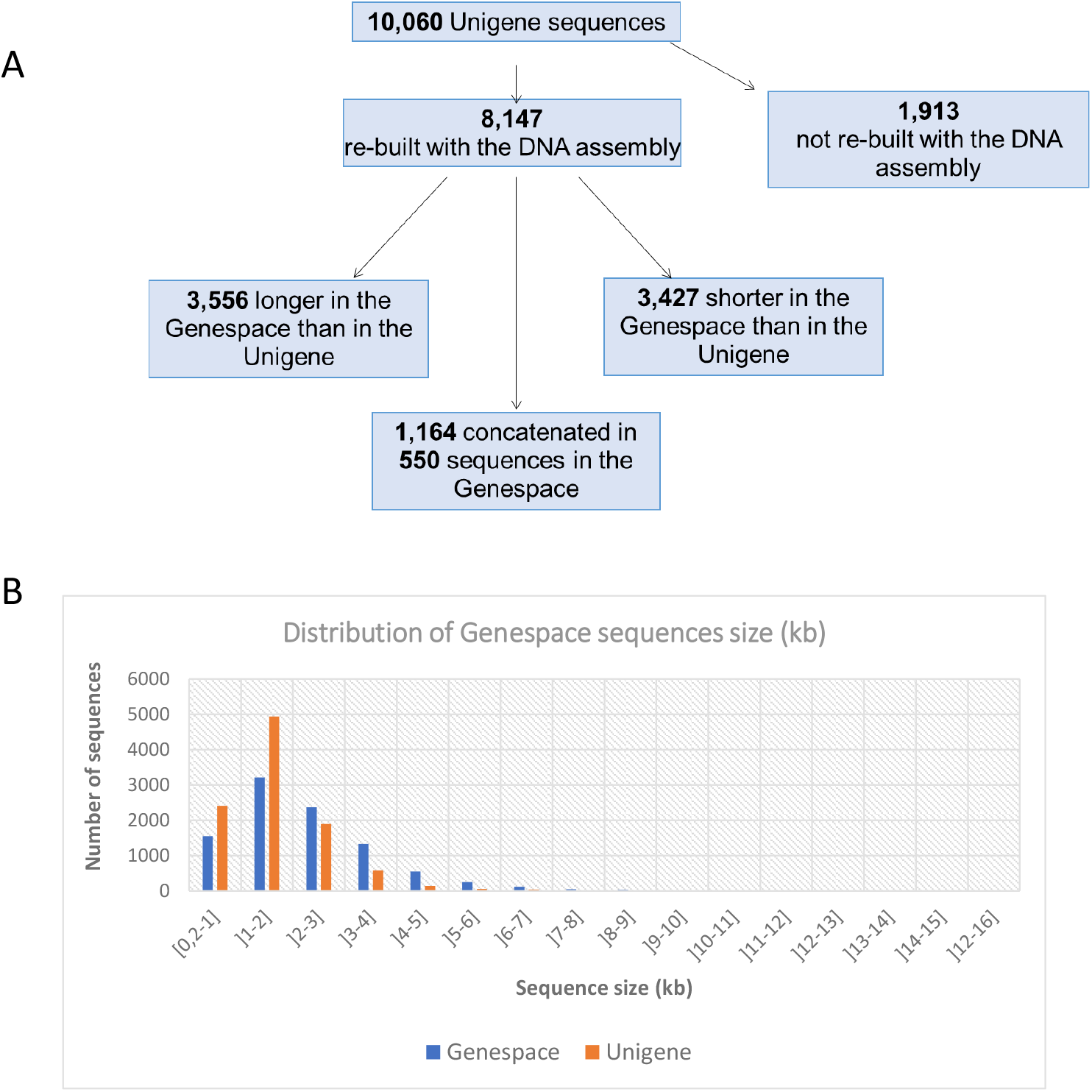
Number and size of sequences built in the Genespace in comparison to the Unigene. **(A)** A BLASTN alignment was perfomed to evaluate the number of sequences from the reference Unigene that were recover in the Genespace at the end of the 8ith iteration of transcriptome-guided DNA assembly. **(B)** In the Genespace, we noticed a decrease in the number of sequences with a size <2kb and an increase in the number of sequences with a size >2kb

**Fig. 4:**
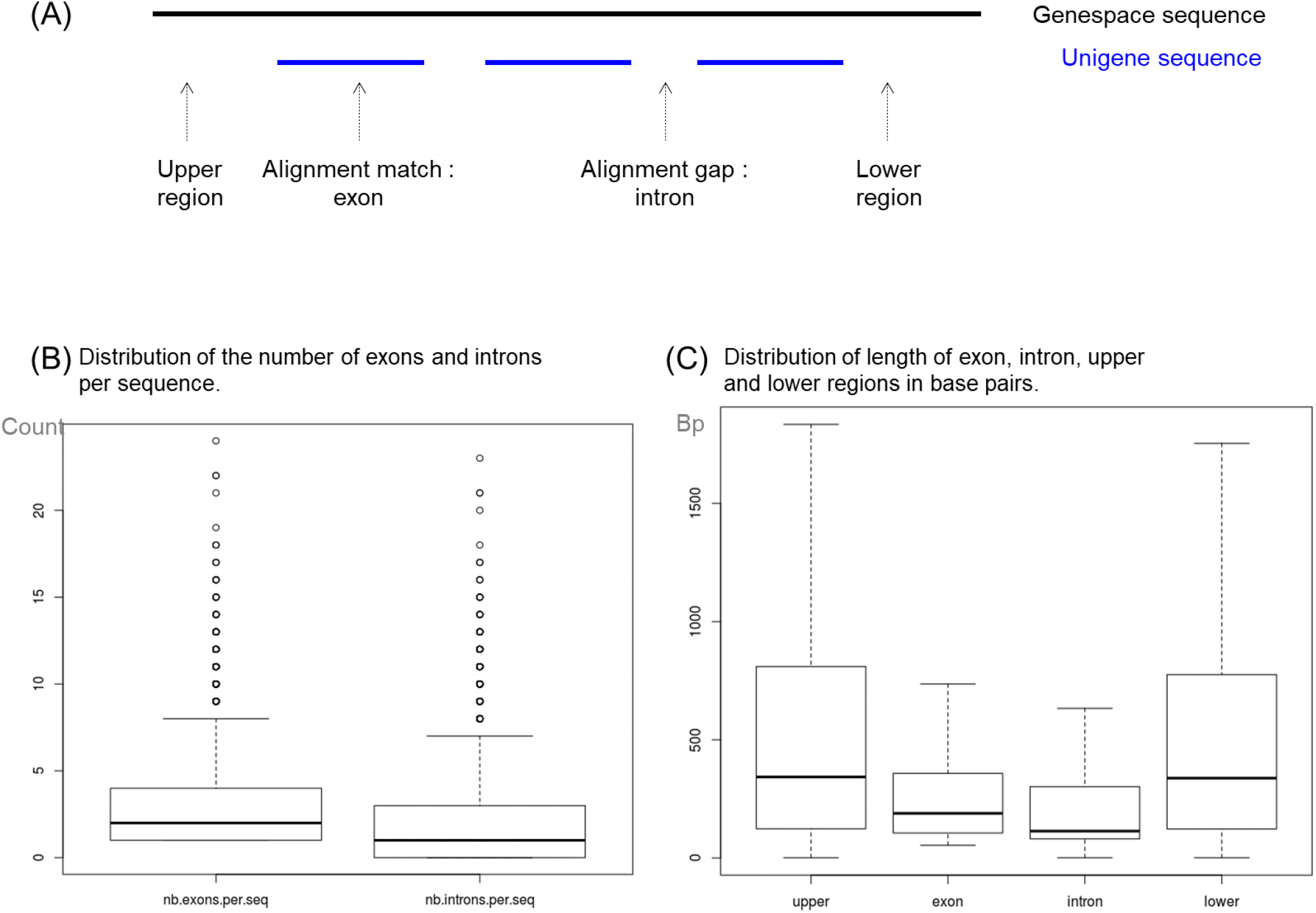
Distribution of exons and intron in Genespace sequences. **(A)** Diagram of the Blastn alignment of an Unigene sequence against its corresponding Genespace sequence. Unigene sequence, built from RNA sequencinfg reads, are mainly compound of exonic regions, whereas Genespace sequences, built from DNA sequencing reads are compound of exonic and non-coding regions. This results in a dashed alignment when aligning both type of sequences, indicative of the position, the number and the length of introns and exons on the Genespace sequences. **(B)** Boxplot of the distribution of the number of exons and introns per Genespace sequence. **(C)** Boxplot of the distribution of the length (in bp) of exon, intron, upper and lower regions on the Genespace sequences. Outliers have been removed from the boxplot for a better readability of the graph.

For SNP detection, we built a reference set called “Genespace” composed of all the sequences initially present in the Unigene but in their longest form either coming from the de novo assembly of RNA-seq, or from the DNA-seq assembly, with the assumption that the longer the sequence, the more introns there are. The sequences not reconstructed with the DNA assembly were also included. The Genespace resulted in a set of 9,446 sequences (21.4 Mb).

### SNP detection

The first step consisted in conducting 16 individual mapping jobs with the DNA sequencing reads from the 16 clones. For each mapping job, almost 6% of the DNA-seq mapped on 85% (more than 8000 sequences) of the Genespace sequences and, at least 90% of the length of each reference sequences was covered by one or more DNA sequencing reads. While removing the 5% extreme values, the mean depth of coverage of the sequences ranged from 9X to 49X. A total of **1,442 Genespace sequences** (1.3Mb – 6% of the Genespace size) were not successfully mapped with DNA-seq reads in none of the 16 mapping jobs. These sequences were part of the 1,913 Unigene sequences that were not rebuilt with the DNA-seq assembly (see previous section). Further investigation indicated that these sequences corresponded to sequences from micro-organisms known to be part of plants rhizosphere (data not shown). Microorganisms DNA was extracted and sequenced simultaneously with the plant DNA. Sequencing was good enough to allow very good sequence assemblies that passed the filters against the SwissProt database and ultimately proved not to be orthologous sequences.

Depending on the individual, a total of 39% to 64% of the SNP identified with CLC were concordant (position + genotype calling) with those identified with GATK. The selection of the subset of concordant SNP between the two tools led to the construction of a genotyping matrix of 16 individuals * 359,323 SNP (Fig 5, S5 Table).

**Fig. 5:**
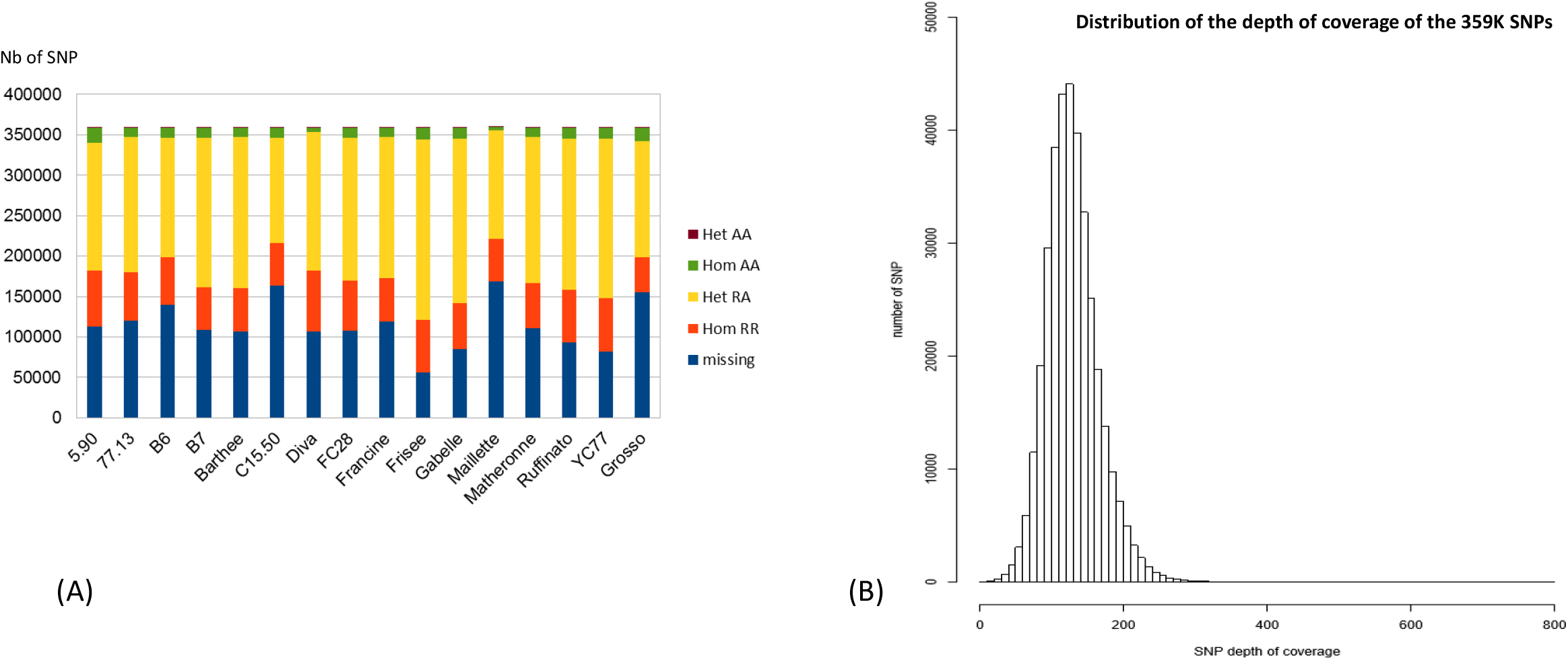
Results fo SNP detection in 16 lavender clones. **(A)** A total of 359K SNP were selected from SNP calling with GATK and CLC. The genotyping information at these sites for the clones studied are represented with barplots.’Het AA’ stands for ‘Heterozygote for two alternate alleles’, ‘Hom AA’ stands for ‘Homozygous for an alternate allele’ Het RA’ stands for ‘Heterozygous with one reference allele and one alternate alleles’, ‘Hom RR’ stands for ‘Homozygous for the reference allele’ and ‘missing’ inidcates no genotyping info due to a low depth of covarage or a low quality for the SNP in that genotype. **(B)** Distribution of the depth of coverage of SNP selected in the 359K gentoyping matrix.

These SNP were spread on 7,332 Genespace sequences with a mean value of 1 SNP per 90 base pairs (bp). Considering the mapping information from the 16 genotypes studied, each polymorphic site conserved in the matrix was supported by 5 to 800 reads, with a median value of 130 reads (Fig 5b). The minimum value of 5X was tolerated when a polymorphic site was detected, or reached sufficient quality, in only one genotype. A total of 61% of the SNP had a minor allele frequency (MAF) ranging from 0.4 (exclude) to 0.5, 29% had a MAF value in the range 0.05-0.4 and 9% had a MAF value inferior or equal to 0.05. The clear majority (98%) of the conserved 359K SNP were biallelic (S5 Table).

On average, within this matrix of 359K SNPs, a clone basically had about 50% of the SNPs at heterozygous state, 20 % at homozygous state and 30% of missing data (Fig 5a, S5 Table). Missing data were mainly due to our stringent settings for SNP calling filtering steps. Interestingly, 114 to 1,281 putative private polymorphisms (*i.e*. allele found in one clone but not the others) were found within the lavender clones, with 20 to 121 SNP being at homozygous state (S5 Table). Moreover, 30,144 putative private SNP including 991 SNP at the homozygous state were identified between the lavandin Grosso and all the lavender clones.

### Genetic distances analysis of 16 lavender clones

The 9,449 polymorphic sites generating no missing data were used to calculate genetic distance between the 16 clones with the Provesti method (S6 Table, S7 Fig). Rare alleles within the genotyping matrix were deliberately preserved in the dataset in order to maximize the discriminations between the clones. The pairwise genetic distance matrix was used to perform a neighbor joining tree with 1000 bootstraps (Fig 6). Some notable clusters were observed. Firstly, and as expected, we observe that ‘Grosso’ appeared to be divergent from the lavender clones. Secondly, some tight clusters were observed: C15.50-B7, Diva-Maillette, Ruffinato-5.90 and Gabelle-Frisée (the two “blue lavender” of the dataset). The bootstrapped trees suggested that the clone 77.13 shared a common ancestor with Ruffinato and 5.90, though 77.13 seems to have more diverged than the two later from the shared ancestor. Finally, it seems that the known geographical origin France / Bulgaria was not structuring in our dataset as B7 was genetically closer to C15.50 than to B6, the other Bulgarian clone.

**Fig. 6:**
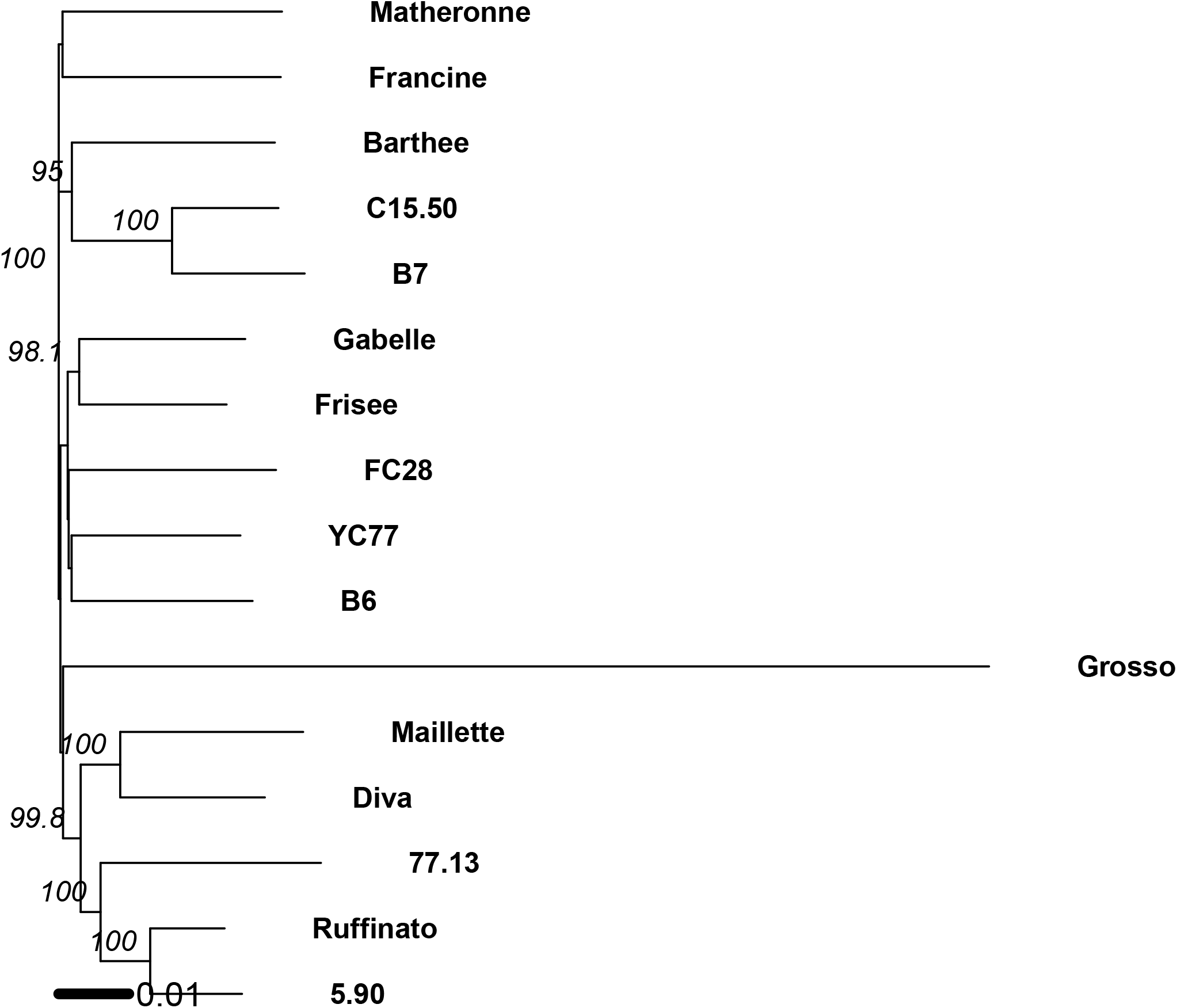
Phylogenetic tree (Neighbour Joining method) of the studied lavender clones.

**Fig. 7:**
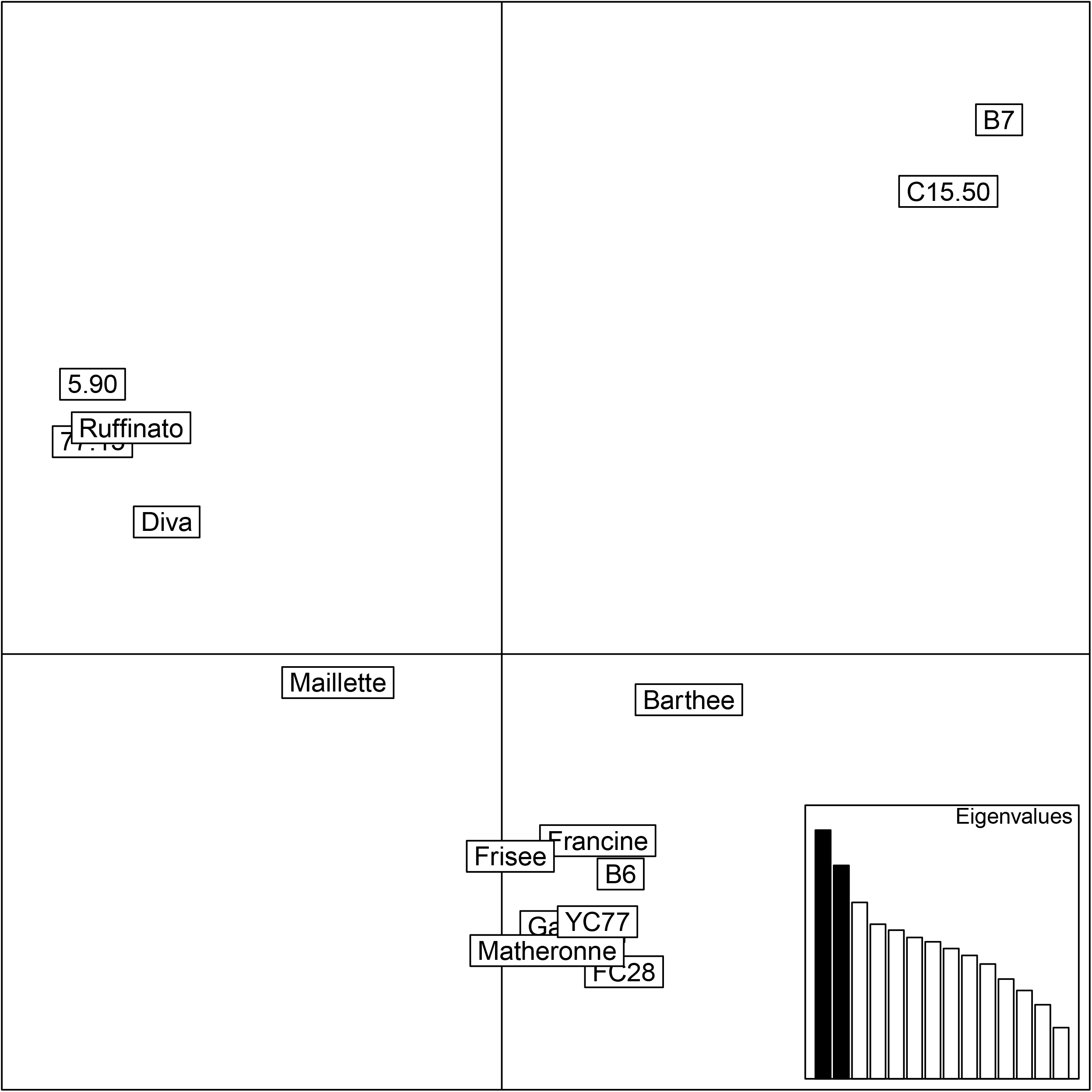
Principal Component Analysis of the genetic distances between the 15 lavenders clones. The PCA was performed with genotyping matrix of the 15 clones and 9449 SNP markers.

Results obtained on the PCA with the 16 clones were analogous to the hierarchical clustering result (S8 Fig). The main axes of the PCA performed with the 16 clones split the lavandin ‘Grosso’ from the lavenders. The second principal axe was due to the group ‘5.90’-‘Ruffinato’-‘77.13’-‘Diva’ and the group ‘C15.50’-‘B7’. The first three axes explained 23.37%, 10.04% and 8.60 %, respectively, of the variability observed between these clones. To better assess the intra-specific diversity, a PCA performed between the lavenders (Fig 6) confirmed that the distinction between ‘C15.50’-‘B7’ and ‘5.90’-‘Ruffinato’ explained most of variability along the first axe. The first three axes explained 13.07%, 11.22% and 9.27%, respectively, of the variability observed between these clones. The clones B6, Francine, Matheronne, FC28, YC77, Frisée and Gabelle appeared to be relatively close to each other’s.

## Discussion

In this study, we proposed a methodology for the rapid development of a large scale molecular resources for an orphan species. We studied as an example *Lavandula angustifolia*, a species of great economic importance from the *Lamiaceae* family and the Medicinal and Aromatic Plant (MAP) sector. Our goals were, firstly to develop genomic reference sequences thanks to *de novo* RNA-seq assembly coupled with a transcriptome-guided DNA-seq assembly. Secondly, we used these resources for SNP mining and finally tested the SNP within the scope of a phylogeny analysis of lavender clones.. In this section, we discuss the efficiency, the advantages and the areas requiring improvement of the method proposed.

### Construction of good-quality gene sequences through coupled use of RNA and DNA sequencing data

We used different licensed and free software for our analyses. However, the method presented in this study for introducing introns can be used with other tools, including using only open source software. The bioinformatics platforms provide users with suite of tools and analyzes to make and validate a *de novo* assembly of RNA-seq such as DRAP [46] or the Trinity protocol [30]. Likewise, the transcriptome-guided DNA-seq assembly protocol can be performed with any tool allowing to map reads to a reference genome and then extract reads that mapped in proper and broken pairs.

#### De novo transcriptome assembly

The quality of the assembly is primary based on the quality and the quantity of sequencing reads. Based on previous studies, in which an amount of 20-40 million of reads [47] originating from various tissues were recommended to get a comprehensive assembly, we could say that we had enough data to get relevant sequences from RNA data. To assess the assembly quality, it has been demonstrated that the best metrics included (i) the proportion of reads mapping back to the assembly, (ii) the recovery of conserved, widely expressed genes and (iii) the total number of Unigene sequences [47,48].

In this study, we used the tool TRANSRATE to calculate the proportion of reads that mapped back to the *de novo* assembled contigs. Results indicated that 94% of reads pairs mapped to the contigs but this value dropped to ~60 % when considering reads that mapped in a “good manner” as defined by Smit-Unna *et al* [31]. It is difficult to compare this value since this metric is not yet widely used. Similar studies only provide the global percent of reads mapping back to the reads – which, in our case, indicate that the assemblies are well supported by reads data. Moreover, we obtained TRANSRATE scores of 0.184 and 0.198 for optimized assemblies obtained with CLC and TRINITY, respectively. This indicated that our assembly was better than 25 % of the published assemblies tested in Smith-Unna *et al*. [31].

To assess the recovery of genes, we chose to perform a BLASTX alignment analysis against the SwissProt database. To build the reference Unigene, we only selected assembled sequences that covered at least 70% of their hit in the database with at least 30% of similarity. With these supplementary selective conditions, we restricted ourselves to a narrow annotatability of the reference Unigene and we may have missed out some well assembled sequences or biased our reference Unigene towards well-conserved sequences between species. However, according to our goal that was to develop molecular resources with a high level of reliability, this choice was appropriate to prevent from keeping sequences with assembly errors, which could lead to poor mapping quality for SNP detection [49,28]. Despite filtering steps and due to the database selected for BLASTX alignments, contaminant sequences were identified in the 10,060 selected sequences of the Unigene and removed to **obtain a cleaned reference set of 8, 640 coding sequences**.

Gathering the results obtained from two or more assemblers allow to reduce the number of sequences due to bioinformatics artefacts and thus, to increase the assembly quality [28]. In our reference Unigene, almost 90 % of selected sequences where assembled with both Trinity and CLC indicating the high-quality and reliability of the Unigene developed herein.

Lastly, to complete the information available for assembled reference sequences, we performed *in silico* functional annotations. Due to the selective steps prior to annotation, we have reach high annotability of our reference Unigene. Annotation information may be relevant for genetic studies based on candidate genes approaches such as association mapping analyses.

#### Introns insertion

To perform a good SNP detection, the first step consisting in the alignment of sequencing reads from individuals on a reference sequence is of great importance. The originality of this study lies on the insertion of intronic sequences within the exons of the Unigene sequences to recover full-length gene sequences. This has the advantages to improve read mapping quality at exon/intron junctions and to increase the number of reads mapping on the reference sequences, **the two resulting in increasing the putative number of SNP detected**. In addition, there is potentially more polymorphism in the introns than in the exons because these regions are less subject to selection pressure. **Thus, by adding introns, we maximized the possibility of finding polymorphism between genetically close individuals**.

In this study, DNA assembly for introns insertion allowed the reconstruction of 81 % of the sequences initially present in the reference Unigene versus 88% in the initial study of Aluome *et al*. [27]. Among these reconstructed sequences, we found that the size of some sequences was shorter in the Genespace than in the Unigene. There are different hypotheses to explain this partial reconstruction. The first one would be that there might be transcripts with introns in the reference Unigene since intron retention is a common splicing mechanism in plant [50,51]. By selecting the longest form of sequences, we might have enriched our reference Unigene with sequences where the introns were retained and thus that could not be improved with the DNA assembly. Another cumulative hypothesis would be related to sequencing depth and the high polymorphism frequency in the clone used in this study. Since the sequencing depth reached in the RNA-seq was larger than in the DNA-seq, some complex sequences may have been built in the Unigene but not in the Genespace due to a lower sequencing depth and thus lead to assembly premature stops.

It would be interesting to conduct a structural annotation of the sequences with an appropriate software, in order to have a finer information on the intron/exon organization of the assembled sequences. However, results obtained herein (gene sequences with a mean of 3 exons and 2 introns) are in concordance with mean values observed in plants. Likewise, the size of intronic and exonic regions is in the order of magnitude of what is observed in perennial plants such as *Arabidospsis lyrata* [50], *Fragaria vesca* [52] or trees [53].

### SNP discovery and genetic distances analysis

We found up to 400K polymorphic sites, depending on the genotype analyzed. SNP detection on these clones revealed a high SNP frequency (mean of 1 SNP per 90 bp) and a high level of heterozygosity (more than 60% of heterozygous SNP per genotype). This result is consistent with the outbreed nature of lavender and its domestication history based on massal selection with clonal propagation. Indeed, similar results have been reported for other outbreed perennial plants with recent domestication history such as ryegrass [54] in comparison to species with inbreeding reproduction mode or with a longer domestication history. In the study of Adal *et.al* [22], where SSR markers were developed for lavender, authors also concluded a high polymorphism frequency (1 SSR/ 2.1kb) in comparison to model crops (as rice (1SSR/3.4 kb), wheat (1SSR/5.4 kb) or soybean (1SSR/7.4 kb)).

This study is the first one to report genetic distance analysis at molecular level between a large number of cultivated clonal lavender varieties. The main conclusions that can be drawn from these results are: (i) the globally homogeneous genetic distances between pairs of clones, related to the allogamous nature of the species, a restricted area of cultivation and the varietal creation method; (ii) as well as an absence of structuring related to the geographical origin [10], but this last result must be moderated because we had only two Bulgarian clones in the study. Some results obtained can be interpreted in light of the information available for the clones studied. In particular, the cluster compound of Gabelle and Frisée can be explained by the fact that these two are “blue” lavender, selected for the persistence of the flowers after the cut, whereas all the other lavenders studied in our collection had been selected for their quality and / or yield of essential oil. Similarly, from an agronomic point of view, the Diva and Maillette lavender are distinguished from other cultivars by their very good yield (number or flower/feet), even if they are distinguished by their chemotype (CRIEPPAM, personal com.). Moreover, the results obtained in this study are consistent with previously published results. Notably, the low genetic differentiation within *L. angustifolia* has been previously reported [28]. In a study including wild and improved lavender populations, blue lavenders and clonal varieties - some of which were included in our collection, Chaisse *et al*. [55] had also reported the genetic proximity between C15.50 and B7 as well as a notable structuring of the diversity between the blue lavenders and the clonal varieties selected for their essential oil content. The other observed clusters cannot be interpreted because of lack of information on these cultivars. Indeed, most of these clones were collected from fields of open-pollinated varieties, and these varieties came from collections in natural environment. Thus, the origin of cultivated clones was lost due to a lack of traceability. However, the results presented here are as many clues to decipher the structuring of genetic diversity within the species and to trace the history of the domestication of *L. angustifolia* and its related species. It would be interesting to carry out a larger scale analysis to explore all genomic resources available for breeding programs in order to build appropriate collections for association mapping studies and to assist breeders in the selection of parental genotypes to initiate breeding programs. Our results suggest that, given the low genetic differentiation between cultivars, high resolution molecular markers such as SNP would be required to accurately explore the genetic diversity of lavender species.

## Conclusion

This study presents an original way to use the RNA-seq and DNA-seq assembly to develop molecular resources on a species for which no genomic information is available, with little bioinformatics resources. This method has the advantage of allowing the detection of SNPs in intronic regions, known to exhibit more variability than exons and thus is also suitable to self-pollinated species or genetically close individuals. The results presented in this study are complementary to those published on lavender since they do not target the terpene biosynthesis pathway and are more exhaustive, thus available for a wider range of applications. Moreover, this is the first reported large-scale SNP development on *Lavandula angustifolia*. All this data provides a rich pool of molecular resource for analyze and preserve the biodiversity of the species in order to optimize breeding strategies.

## Data availability

All sequencing and assembly data generated in the GENOPARFUM project have been deposited at DDBJ/ENA/GenBank under the BioProject number PRJNA391145.

## Supporting information

**S1 Table: List of lavender clones studied**.

(merged PDF)

**S2 Table: RNA and DNA sequencing metrics**.

(merged PDF)

**S3 Table: Reference Unigene annotation results with Trinotate, Trapid and KAAS tools**.

(XLSX)

**S4 Fig.: Distribution of top best blast hit species. Sequences resulting from CLC and TRINITY de novo assemblies were aligned against SwissProt database**.

(merged PDF)

**S5 Table: Detailed results of SNP detection with GATK and CLC (concordant SNP only) between the lavender clones**.

(merged PDF)

**S6 Table: Dissimilarity matrix (provesti’s distances) calculated between the lavender clones**.

(merged PDF)

**S7 Fig.: (A) Boxplot of 9449 SNP depth of coverage. (B) Boxplot of 9449 SNP minor allele frequencies**.

(PDF)

**S8 Fig.: Principal Component Analysis of the genetic distances between the 16 lavenders clones**.

(PDF)

**S9 File: Bash script used for introns insertion in unigene sequences**.

(TXT)

## Acknowledgement

We would like to thank Vegepolys Innovation, more specifically Fabienne Mathis, for their contribution in raising the GenoParfum project. We are grateful to the genotoul bioinformatics platform Toulouse Midi-Pyrenees (Bioinfo Genotoul) for providing computing and storage resources. We would like to thank all our colleagues from INRA (Angers, Orleans, Orsay, Rennes, Versailles) for the enriching and constructive exchanges around this project, especially Patricia Faivre-Rampant for reviewing the article. The authors thank the Ministere of Agriculture for the funding of the Casdar project GENOPARFUM that allowed to perform this study.

## Authors contribution

Conceptualization: JPBB, DB, PG

DNA/RNA extraction and sequencing: AB, JBR, MCLP, BFF

Bioinformatics / data analyses: BFF

Writing: BFF, DB

